# Investigation of mitochondrial phenotypes in motor neurons derived by direct conversion of fibroblasts from familial ALS subjects

**DOI:** 10.1101/2025.02.13.637962

**Authors:** Evan Woo, Faiza Tasnim, Hibiki Kawamata, Giovanni Manfredi, Csaba Konrad

## Abstract

Amyotrophic lateral sclerosis (ALS) is a progressive neurodegenerative disease of motor neurons, leading to fatal muscle paralysis. Familial forms of ALS (fALS) account for approximately 10% of cases and are associated with mutations in numerous genes. Alterations of mitochondrial functions have been proposed to contribute to disease pathogenesis. Here, we employed a direct conversion (DC) technique to generate induced motor neurons (iMN) from skin fibroblasts to investigate mitochondrial phenotypes in a patient-derived disease relevant cell culture system. We converted 7 control fibroblast lines and 17 lines harboring the following fALS mutations, SOD1^A4V^, TDP-43^N352S^, FUS^R521G^, CHCHD10^R15L^, and C9orf72 repeat expansion. We developed new machine learning approaches to identify iMN, analyze their mitochondrial function, and follow their fate longitudinally. Mitochondrial and energetic abnormalities were observed, but not all fALS iMN lines exhibited the same alterations. SOD1^A4V^, C9orf72, and TDP-43^N352S^ iMN had increased mitochondrial membrane potential, while in CHCHD10^R15L^ cells membrane potential was decreased. TDP-43^N352S^ iMN displayed changes in mitochondrial morphology and increased motility. SOD1^A4V^, TDP-43^N352S^, and CHCHD10^R15L^ iMN had increased oxygen consumption rates and altered extracellular acidification rates, reflecting a hypermetabolic state similar to the one described in sporadic ALS fibroblasts. FUS^R521G^ mutants had decreased ATP/ADP ratio, suggesting impaired energy metabolism. We then tested the viability of iMN and found decreases in survival in SOD1^A4V^, C9orf72, and FUS^R521G^, which were corrected by small molecules that target mitochondrial stress. Together, our findings reinforce the role of mitochondrial dysfunction in ALS and indicate that fibroblast-derived iMN may be useful to study fALS metabolic alterations. Strengths of the DC iMN approach include low cost, speed of transformation, and the preservation of epigenetic modifications. However, further refinement of the fibroblasts DC iMN technique is still needed to improve transformation efficiency, reproducibility, the relatively short lifespan of iMN, and the senescence of the parental fibroblasts.

## Introduction

Amyotrophic lateral sclerosis (ALS) is a debilitating neurodegenerative disease marked by the progressive degeneration of upper and lower motor neurons, leading to paralysis and eventual death within 3-5 years after diagnosis. Approximately 10% of ALS cases are familial (fALS), caused by identifiable genetic mutations in genes including SOD1, C9ORF72, TARDBP, FUS, and CHCHD10. These mutations cause a range of molecular alterations that contribute to disease pathogenesis. Pathogenic mechanisms encompass diverse cellular processes such as RNA metabolism, protein homeostasis, neuroinflammation, oxidative stress, and mitochondrial dysfunction. Understanding these processes is crucial for developing effective therapies, which are still lacking. Animal models and cell lines have provided many insights in disease mechanisms and are routinely used to screen drugs and test new therapies. However, they have limitations in recapitulating the human and neuronal specific aspects of the disease. To address this problem, technologies to generate motor neurons from human cells, such as pluripotent stem cells (iPSC), have been developed and have provided valuable culture human neuron models for studying ALS mechanisms. In addition to iPSC, more recently other approaches to generate human motor neurons are being developed. Notably, direct conversion (DC) of fibroblasts to induced motor neurons (iMN) without the need for epigenetic reprogramming steps, might offer a complementary in vitro approach to iPSC for studying neurodegenerative diseases^1,2^, including ALS^3–5^.

DC of fALS fibroblasts has been used as a platform for drug screening and for mechanistic studies^6,7^. Previous studies utilizing fALS iMN generated via DC showed phenotypic differences compared to controls. For example, various FUS mutant lines showed mislocalization of FUS protein into stress granules and reduced iMN viability ^8,9^. Moreover, C9orf72 mutant DC iMN showed presence of RNA foci and repeat associated non-ATG translation yielding dipeptide repeat proteins^10^. In addition, cytoplasmic TDP-43 mislocalization was detected in TDP-43 mutant iMN, but not in their parent fibroblasts^11^. These studies suggest that DC iMN can provide a viable human iMN platform for studying disease mechanisms and testing therapeutic approaches. However, although mitochondrial dysfunction and energy metabolism alterations are thought to play an important pathogenic role in ALS^12,13^, it has not yet been studied in fALS DC iMN.

In this study, we generated DC iMN from fibroblasts of individuals affected by various forms of fALS to investigate mitochondrial phenotypes. We also compared efficacy of DC among multiple independent cell lines harbouring different fALS mutations and addressed iMN viability under baseline culture conditions and in responses to small molecules that modulate mitochondrial function and oxidative stress. Importantly, we found that DC efficiency varies substantially among fibroblast lines, independent of mutation, passage number, and donor’s age. We developed a machine learning algorithm to identify and follow longitudinally DC iMN and to perform mitochondrial functional and morphological measurements. Despite inter-line variability, in a subset of fALS mutants, we were able to detect mitochondrial and cell viability phenotypes. The latter were largely rescued by treatment with small molecules. These findings suggest that DC iMN may serve as a viable platform to investigate metabolic alterations in fALS.

## Materials and Methods

### Skin biopsy, fibroblast cultures and iMN differentiation

After informed consent, a skin biopsy was obtained from the volar part of the forearm. Skin biopsies were de-identified to protect subject’s identity. Fibroblast samples were provided to our laboratory as deidentified, coded samples. Some lines were obtained from the NINDS catalogue of motor neuron disease fibroblasts. Skin fibroblasts were cultured as described previously^14–18^ in Dulbecco modified Eagle medium (DMEM) (ThermoFisher Scientific, Waltham, MA) supplemented with 25 mM glucose, 4 mM glutamine, 1 mM pyruvate, and 10% foetal bovine serum (hereafter growth medium). All cultured fibroblast lines were regularly tested for mycoplasma infection. Lines were subjected to DC at passages ranging between 5 and 12. We did not observed loss of contact inhibition in any of the lines.

Fibroblasts were transformed into iNMs based on the protocol of the Yoo group^19^ with minor modifications. In brief, fibroblasts were infected with the following lentiviral constructs, pTight-9-124-BclxL (Addgene #60857), rtTA-N144 (Addgene #66810), and pCSC-ISL1-T2A-LHX3 (Addgene #90215), at ∼1×10^6 TU/ml / 300 000 cells/well in 6 well plates on day 0 (D0) in the presence of polybrene (Sigma, 6 µg/ml) and using spinfection (900g for 90mins @37°C). Cells were then differentiated as described^19^. On D6, cells were replated in 96 well glass-bottom imaging plates (P96-1.5H-N, Cellvis) for viability tracking, immunofluorescence, and TMRM live imaging or in 18 well µ-slide glass bottom slides (ibidi, Cat.#:81817) for live imaging of Perceval and kymograph imaging, at 25 000 cells/well. Antibiotic selection by 3 µg/ml Puromycin was from D11-14.

### Viability tracking

On D8 cells were transduced with an eGFP expressing lentiviral vector under the synapsin promoter, pHR-hSyn-EGFP (Addgene #114215), at 0.5×10^6 TU total / well without polybrene or spinfection. Wells were imaged starting at D12 until D46 every second day using an ImageXpress Pico high-content imaging system (Molecular Devices) at 10x magnification (0.69µm/pixel). A YOLOv8 model was fine-tuned to detect cells exhibiting neuronal morphology (small bright round cell body with neurites) over time, and the built-in BoT-SORT algorithm was used to track individual iMN. Survival analysis and COX hazards modelling was done using tracks of 500 randomly selected iMN cells / fALS or control lines using the lifelines python package.

### Neurite morphology assessment

The iMN cultures were imaged at 20x magnification (0.345µm/pixel) on D30 for neurite morphology assessment. The image analysis pipeline is illustrated on Fig. 2A. Images were background-subtracted, and contrast was adjusted in ImageJ. Cell bodies were detected by gaussian blurring and thresholding, followed by size exclusion. The cell body and thresholded images were processed in CellProfiler to generate single-cell segmentations based on the cell bodies using the watershed algorithm. Cropped cell bodies were also used to classify single cells as either iMN, non-iMN, or dead cells by a ResNet-50 deep learning model trained for this task. Single cell iMN segmentations were then individually skeletonized and measured in ImageJ.

### Immunofluorescence

iMN were differentiated as described above, but without pHR-hSyn-EGFP transfection. On D30 cells were fixed for 30 minutes in 4% paraformaldehyde, blocked by BSA in PBS+20 mM glycine, incubated over night with anti TDP-43 rabbit antibody (Proteintech, cat 10782-2-AP, 1:200), at 4°C, followed by washing in PBS+20 mM glycine and staining with Cy5 anti-rabbit secondary antibody (Jackson, 1:500), and Hoechst 33342 (1µg/ml). Images were taken using an ImageXpress Pico high-content imaging system (Molecular Devices) at 40x magnification (0.1725µm/pixel). The image analysis pipeline is illustrated on Fig. 3A. Images were background subtracted, and contrast was adjusted in ImageJ. Enhanced contrast images were created to aid single cell segmentation by equalizing histograms (EQ) of the TDP-43 and Hoechst channels and summing the two images pixelwise (Fig. 3 B). This enhances background signal in the cell bodies and neurites. Nuclei were segmented by gaussian blurring, thresholding, followed by size exclusion. The nuclei and EQ image was processed in CellProfiler to generate single-cell segmentations. The nuclei were also used to cut out cell bodies to be used to classify single cells as either iMN, non-iMN, or dead by a ResNet-50 deep learning model trained for this task. Single-cell segmentations were then paired up with filtered iMN nuclei segmentations in imageJ and TDP-43 intensities were measured in both the nucleus and the cytosol.

### Mitochondrial membrane potential and morphology

On D30 of differentiation iMN cultures were stained by Hoechst 33342 (1µg/ml), and TMRM (15nM) for 45 minutes. Images were taken on an ImageXpress pico (Molecular devices) at 40x magnification (pixel size = 0.1725 µm). Single-cell segmentation and iMN filtering was done on the TMRM images, essentially as described above for TDP-43 immunofluorescence. Individual iMN were then analysed in ImageJ. For each neuron individual mitochondria were segmented and measured. Segmented mitochondria were also skeletonized and measured for size and shape. Single mitochondria measurements were then averaged for each cell.

### Mitochondrial motility

Cells were seeded on 18 well glass bottom (ibidi Cat.No:81817). On D30, cells were stained by TMRM as described above. Live imaging was done using a DMIRB Inverted microscope (Leica), equipped with temperature, humidity, and gas control. iMN were imaged at 40x for 10 minutes at 0.33Hz. Neurites were identified manually in ImageJ to generate kymographs of individual mitochondria. Proportion of moving and stationary mitochondria were assessed. For motile mitochondria, antero- and retrograde travel distances, average speeds, and maximum speeds were measured.

### ATD/ADP ratio measurements by PercevalHR

Differentiations were done as described above for mitochondrial motility. Cells were infected with lentivirus for the biosensor PercevalHR^20^ on D26. On D30, media was changed to Krebs-Ringer Bicarbonate Buffer supplemented with 1mM Ca^2+^ and 5 mM glucose. Live imaging was done on the Leica DMIRB Inverted microscope with binning set to 4. iMN were imaged at 20x for 25 minutes at 0.2Hz. Oligomycin (1 µg/ml) was added at 5 minutes and 2-deoxy-D-glucose (25 mM) at 15 minutes. iMN were identified and segmented manually and blue (405 nm) and green (490 nm) fluorescence signals were quantified to calculate the 490/405nm ratio, as described^20^.

### Measurements of oxygen consumption and extracellular acidification

Oxygen consumption rate (OCR) and extracellular acidification rate (ECAR) were measured with a XF96 Extracellular Flux Analyzer (Agilent, Santa Clara, CA). Cell lines were seeded in 6 replicate wells of a XF 96-well cell culture microplate (Agilent). On D30, the medium was replaced with 200 μL of XF Assay Medium (Agilent) supplemented with 5 mM glucose, 1 mM pyruvate and 4 mM glutamine, pre-warmed at 37°C, iMN were degassed for 1 h before starting the assay procedure, in a non-CO_2_ incubator. OCR and ECAR were recorded at baseline followed by sequential additions of oligomycin (1 µg/ml), FCCP (2 µM), and Antimycin A (0.5 µM) plus Rotenone (0.5 µM) and 2-deoxy-D-glucose (25 mM). Non-mitochondrial oxygen consumption and non-glucose driven acidification (rate values after all inhibitors added) were subtracted from all OCR and ECAR values respectively, and technical replicates with negative values were discarded for both ECAR and OCR. Values were normalized by the number of cells in each well, measured by imaging of nuclei (Hoechst) and cytosol (calcein) fluorescence imaging.

### Statistical analyses and code availability

For most measurements, we used one-way ANOVA (Graphpad prism), with p-value correction by Dunett’s multiple comparisons test, to compare ALS mutants to controls. For survival analyses the python lifelines package was used to fit Kaplan-Meier survival curves and test differences using the pairwise logrank test. Hazard ratios and confidence intervals were calculated using Cox’s proportional hazard model. Scripts and readme files for usage are available on out github page (https://github.com/csabak/ALS_DC_iMN).

## Results

### DC iMN efficiency varies among fibroblast lines independent of genotype

To assess the feasibility of miR-9/9*-124 plus ISL1/LHX3 based DC in our laboratory, we used a control fibroblast line from a 64-year-old individual. Fibroblasts were efficiently transduced by lentivirus, as evidenced by immunofluorescence for nuclear ISL1 three days after ISL1/LHX3 lentiviral infection (Fig. 1A). The puromycin selection was effective, as untransfected fibroblasts showed 100% cell death (not shown) after three days in the presence of 0.5 µg/ml puromycin (in contrast with 3 µg/ml puromycin for 10 days during conversion). Upon transformation, over time an increasing proportion of cells developed small round cell bodies and neurite-like structures, as evidenced by brightfield microscopy (Fig. 1B). Cells did not divide during the fibroblast to iMN DC, and most cells remained stationary regardless of morphology. On D30, iMN showed strong protein expression of the neuronal markers NeuN, MAP2, Tuj1, and Syntaxin, as well as cholinergic markers, such as CHAT, and motor neuron (MN) specific markers, such as Lim3, and Mnx1 (Fig. 1C). Bulk mRNA expression analysis of markers confirmed the expected decrease in fibroblast specific markers, such as VIM and CAV1, and the increase of neuronal and MN-specific markers (Fig. 1D).

**Figure 1.**
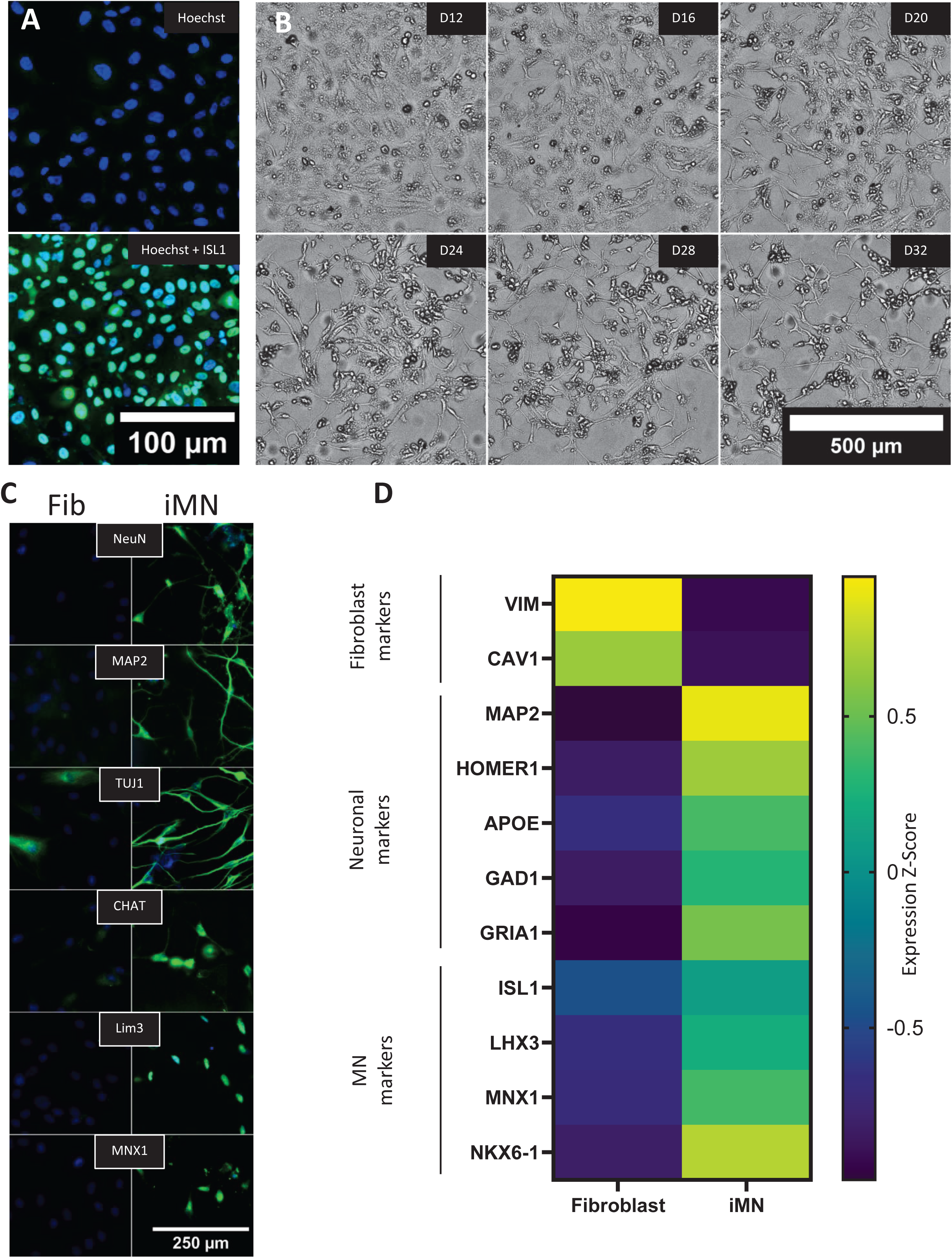
Characterization of iMN directly converted from fibroblasts. (A) Immunofluorescence images of fibroblasts before and after lentiviral infection with ISL1 and LHX3, showing nuclear expression of ISL1 in fibroblasts after three days of infection. Scale bar, 100 µm. (B) Brightfield microscopy example of morphological changes during direct conversion from fibroblasts to iMN between D12-D32. Cells develop neuron-like morphology with rounded cell bodies and neurite extensions. Scale bar, 500 µm. (C) Immunofluorescence staining for neuronal (NeuN, MAP2, TUJ1) and motor neuron-specific markers (CHAT, Lim3, MNX1) in fibroblasts (Fib) and iMN. Staining demonstrates expression of neuronal markers in iMN on day 30 of differentiation. Scale bar, 250 µm. (D) Heatmap of expression Z-scores for fibroblast, neuronal, and motor neuron-specific markers in fibroblasts and iMN. Fibroblast-specific markers (VIM, CAV1) are downregulated, while neuronal and motor neuron markers (MAP2, HOMER1, APOE, GAD1, GRIA1, ISL1, LHX3, MNX1, NKX6-1) are upregulated in iMN (n=24).

Differentiated cultures consisted of cells with variable morphologies, and a subset of cells expressed ISL1 and Lim3, indicating a MN lineage (Fig. 2A). We tested several antibodies, but did not find any marker that could distinguish cells with neuronal morphology, even though all antibodies exhibited significantly lower reactivity on fibroblasts (Fig. 1C), suggesting that upon transduction most cells changed gene expression profiles, but only a portion acquired neuronal morphology.

**Figure 2.**
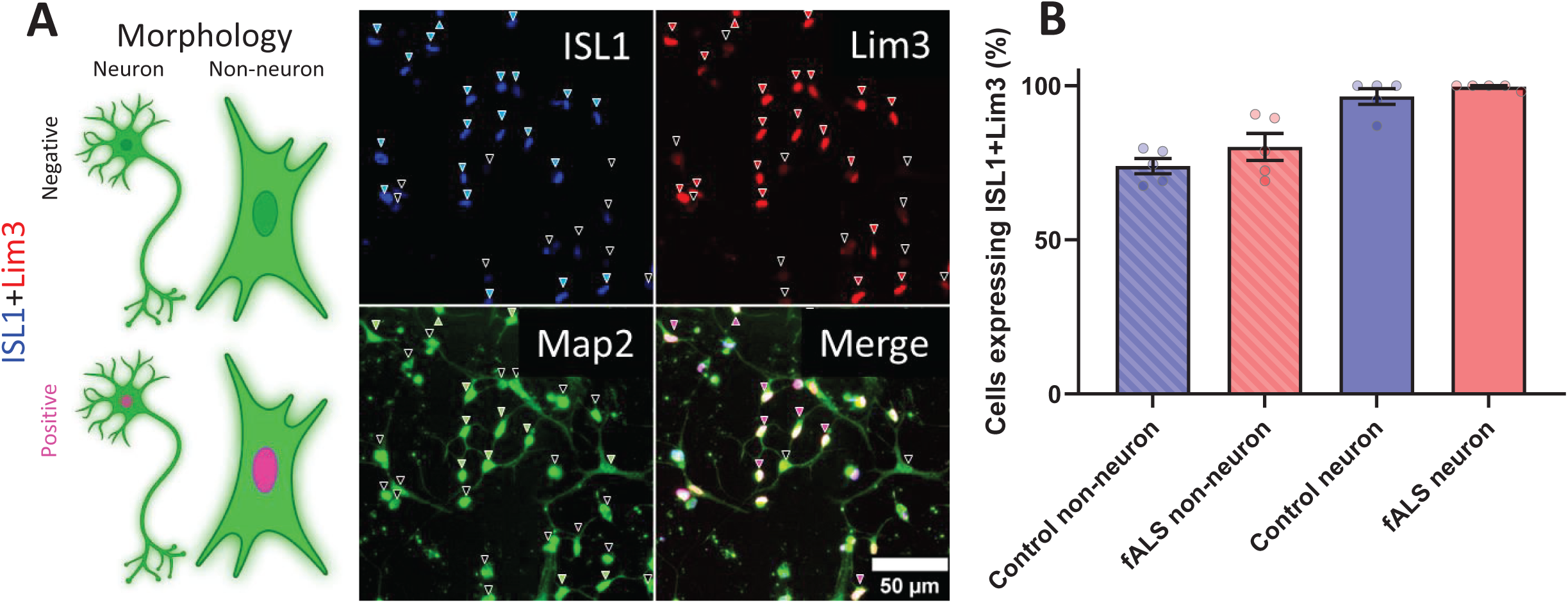
Expression of MN markers ISL1 and Lim3 in control and fALS-derived iMN cultures. (A) cartoon and example images showing cells expressing or lacking motor neuron markers and neuronal morphology. The immunofluorescence images show cells expressing neuronal marker MAP2 (green), motor neuron markers ISL1 (blue), and Lim3 (red) in control iMN on Day 30 of differentiation. Arrowheads indicate cells expressing ISL1(blue) and Lim3 (red), or the lack thereof (black on the corresponding panel). Green arrowheads indicate cells on the MAP2 staining exhibiting neuronal morphology, while black arrows indicate non-neuronal or dead morphology. Pink arrowheads on the merged panel indicate iMN (neuronal morphology with ISL1 and Lim3 expression), black arrowheads indicate iNs (neuronal morphology only) on the merged image. Scale bar, 50 µm. (B) Quantification of cells expressing ISL1 and Lim3 across different categories: control and fALS non-neuronal cells, and control and fALS cells with neuronal morphology. The majority of cells with neuronal morphology in both control and fALS cultures exhibit ISL1 and Lim3 expression, confirming successful motor neuron conversion. Data are presented as mean ± SEM, n=5 Control and 5 fALS, for each data point neuron counts ranged 11-55 cells and non-neuron counts 38-110 cells.

Next, we converted 5 control and 5 fALS (1 SOD1^A4V^, 2 C9orf72, 1TDP-43^N352S^, 1 FUS^R521G^) fibroblast lines and immunostained them for ISL1, Lim3 and MAP2 at Day 30 of differentiation to assess the proportion of iMN. Virtually all cells expressed MAP2, most cells expressed ISL1 (Fig. 2B blue arrows), and most ISL1 expressing cells also expressed Lim3 (Fig. 2B red arrows). We counted cells with neuronal morphology based on MAP2 staining (Fig. 2B green arrows) and measured the proportion of neuron-like cells expressing ISL1 and Lim3 (iMN, Fig. 2B purple arrows). Cells classified as non-neuronal based on MAP2 morphology co-expressed ISL1 and Lim3 similarly (76-79% of all cells), with very low variability among lines (Fig. 2C). On the other hand, cells exhibiting neuronal morphology by MAP2 labelling were almost 100% iMN by Lim3 and ISL1 staining, regardless of genotype (Fig. 2C). This finding suggests that there is variability in differentiation efficiency among cell lines and neuronal markers alone are insufficient to identify neurons. Therefore, morphological criteria are needed to correctly classify iMN for single cell analyses.

To compare conversion efficiency and reproducibility across fibroblast lines and to investigate cellular phenotypes, we generated iMN from fibroblast of 7 control, and 17 fALS subjects. The fALS lines harboured SOD1^A4V^ (n=3), TDP-43^N352S^ (n=4), FUS^R521G^ (n=4), and CHCHD10^R15L^ (n=3) amino acid mutations, as well as C9orf72 intronic repeat expansions (n=4). We measured transformation efficiency in individual batches for each of the 24 lines, using morphological criteria on DC iMN transduced with a lentiviral human synapsin promoter eGFP (hSyn-eGFP). We choose this method for efficiency quantification instead of ICC, because the cells with neuronal morphology are sensitive to washing away during the protocol causing an underestimation of iNM. Six days after infection, nearly all cells in our iMN cultures showed green fluorescence, regardless of morphology, which was maintained throughout the differentiation process (Fig. 3A). We assessed differentiation efficiency on D30. We first identified cell bodies in ImageJ by filtering for size and roundness. We then generated training data for a machine learning algorithm to classify centre-cropped images of individual cells bodies as iMN, non-neuron, or dead (Fig. 3B and Supplementary Fig. S1). We found that different donor lines exhibited variable transformation efficiency into iMN, ranging from approximately 5% to 60%. On the other hand, in most cases, transformation efficiency was quite consistent within the same line across different batches (Fig. 3C). This finding suggests that fibroblast cultures from different donors and across different batches have variable degree of differentiation efficiency,. It also suggests that, when studying iMN, it is advisable to utilize and average multiple lines for each mutation. Aggregating the data by genotype, we found that mutants did not exhibit significantly different transformation efficiencies compared to controls (Fig. 3D).

**Figure 3.**
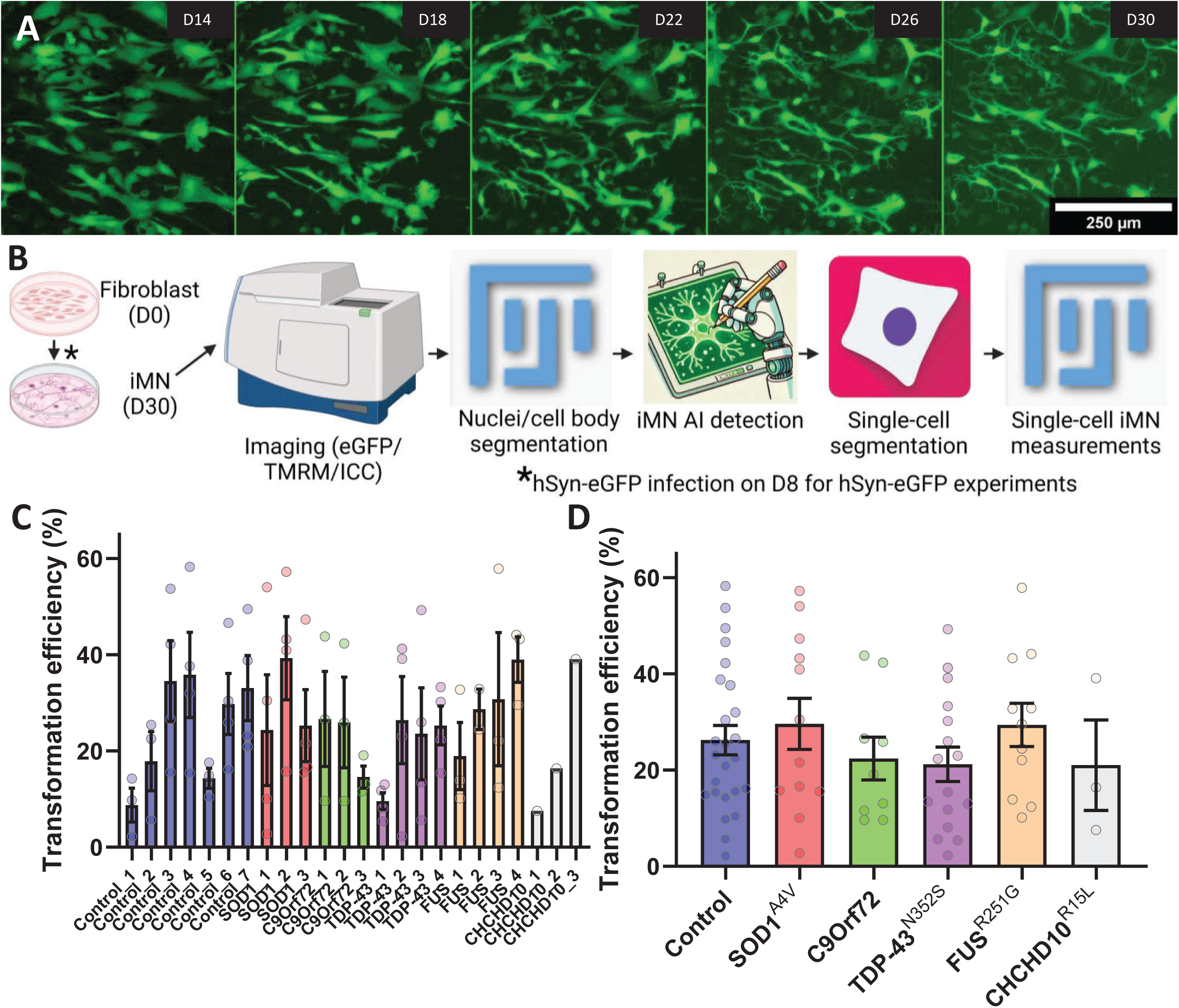
iMN transformation efficiency and analysis workflow for control and fALSpatient-derived fibroblasts. (A) Representative images of hSyn-eGFP expressing iMN cultures over time, showing morphological progression from Day 14 (D14) to Day 30 (D30) of the conversion. iMN exhibit neuronal morphology with well-defined neurite extensions by D30. Scale bar, 250 µm. (B) Schematic of the image analysis pipeline for tracking iMN transformation efficiency and single-cell measurements. The process includes DC, staining and imaging of iMN, nuclei and cell body segmentation, AI-driven iMN classification, and single-cell segmentation for subsequent analyses. hSyn-eGFP infection is introduced on Day 8 (D8) for experiments involving live fluorescent labeling of iMN. (C) Transformation efficiency of fibroblast lines into iMN. Bar graph showing the transformation efficiency (%) of fibroblasts from control and fALS lines into iMN. Each bar represents a different fibroblast line, with control lines shown on the left and fALS lines, categorized by mutation (SOD1^A4V^, C9orf72, TDP-43^N352S^, FUS^R521G^, and CHCHD10^R15L^), shown on the right. Each data point represents an individual batch (n=1-4 per line), and error bars indicate SEM. Transformation efficiency was assessed by measuring the percentage of cells exhibiting neuronal morphology. (D) Transformation efficiency of control and fALS-derived fibroblast lines pooled and expressed as the percentage of cells exhibiting iMN morphology on D30. Error bars represent SEM, n=1-4 for each line (7 control, 3 SOD1^A4V^ 3 C9orf72, 4 TDP-43^N352S^, 4 FUS^R521G^, 3 CHCHD10^R15L^), *p ≤ 0.05, **p ≤ 0.01, ***p ≤ 0.001, ****p ≤ 0.0001.

At D30, we assessed neurite morphology of iMN by analysing hSyn-eGFP images, with additional processing steps to generate skeletons of individual cells that were classified as iMN by the Resnet50 model (Fig. 4A). TDP-43^N352S^ and CHCHD10^R15L^ lines had similar neurite morphology as controls, whereas SOD1^A4V^ iMN had decreased total neurite length per cell (Fig. 4B), and both SOD1^A4V^ and FUS^R521G^ had decreased branching of neurites (Fig. 4C).

**Figure 4.**
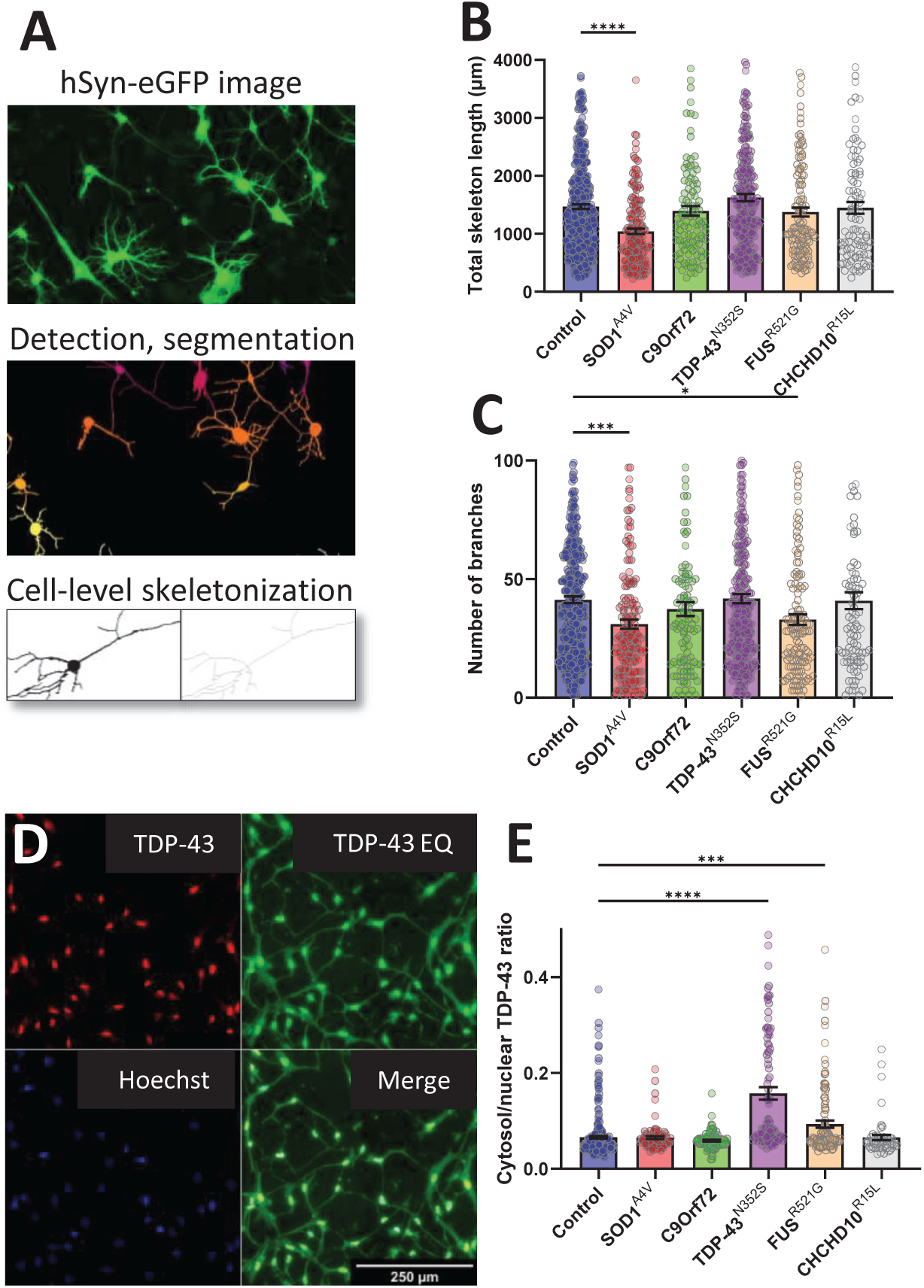
Morphological and TDP-43 localization analysis in iMN from control and fALS subject-derived fibroblasts. (A) Steps of applying the pipeline depicted on Fig. 3B for images of iMN infected with hSyn-eGFP for cell-level skeletalization to assess neurite length and branching complexity. (B) Quantification of total neurite length per cell across control and fALS genotypes, showing significantly reduced neurite length in SOD1^A4V^ iMN compared to controls. (C) Analysis of neurite branch count per cell, with SOD1^A4V^ and FUS^R521G^ mutants showing significantly fewer branches than control iMN. (D) Immunofluorescence staining for TDP-43 and Hoechst in iMN to evaluate TDP-43 cytoplasmic mislocalization, with equalized contrast (EQ) to enhance visualization and segmentation. Scale bar, 250 µm. (E) Quantification of the cytoplasmic-to-nuclear TDP-43 intensity ratio, highlighting increased cytoplasmic mislocalization in TDP-43^N352S^ and FUS^R521G^ mutant iMN compared to controls, whereas other genotypes did not show significant mislocalization. Error bars represent SEM, n=200 cells for each line (6 control, 2 SOD1^A4V^ 2 C9orf72, 2 TDP-43^N352S^, 3 FUS^R521G^, 2 CHCHD10^R15L^), *p ≤ 0.05, **p ≤ 0.01, ***p ≤ 0.001, ****p ≤ 0.0001.

Lastly, since TDP-43 mislocalization is a common feature of ALS postmortem CNS and iPSC models of the disease, as part of the characterization of DC iMN we assessed TDP-43 intracellular localization by immunostaining. Data collection and analysis pipeline were similar to neurite morphology (Fig. 3B) except that Hoechst-stained nuclei, instead of whole cell bodies, were used to detect and centre-crop individual cells, and enhanced contrast images of TDP-43 staining were used for cell segmentation by equalizing the histograms in ImageJ. Equalizing TDP-43 intensity histogram enhanced cytosolic staining and allowed for morphological classification and segmentation of individual cells (Fig. 4D). We found that TDP-43 cytosolic/nuclear ratio was higher in TDP-43^N352S^ and FUS^R521G^ iMN than in controls (Fig. 4E), but not in SOD1^A4V^ and CHCHD10^R15L^ iMN. We did not detect TDP-43 cytosolic inclusions in any of the mutant lines.

### fALS iMN show alterations in mitochondrial morphology and function

To assess mitochondrial membrane potential and morphology we performed live imaging of D30 iMN stained with TMRM (Fig. 5A). Individual iMN were segmented based on morphological criteria using Hoechst-stained nuclei to detect and centre-crop individual cells and enhanced contrast images. We found that whole cell TMRM fluorescence was higher in SOD1^A4V^, C9orf72, and TDP-43^N352S^ iMN compared to controls, unchanged in FUS^R521G^, and lower in CHCHD10^R15L^ iMN (Fig. 5B). TDP-43^N352S^ iMN showed significant differences compared to controls in mitochondrial morphological parameters, including higher number of mitochondria per cell (Fig. 5C), lower average mitochondrial area (Fig. 5D), which were associated with higher TMRM intensity of individual segmented mitochondria (Fig. 5E).

**Figure 5.**
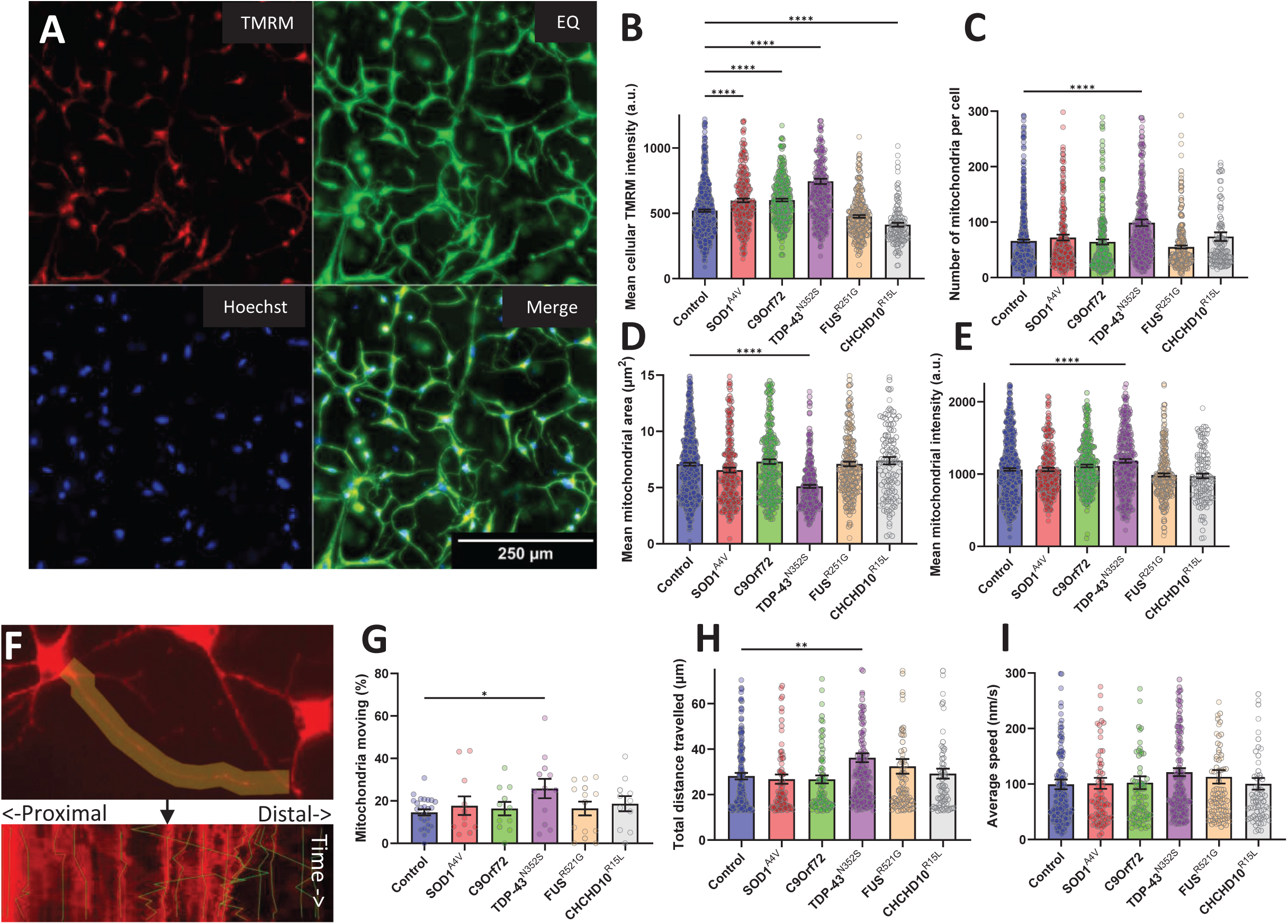
Analysis of mitochondrial membrane potential, morphology, and motility in iMN from control and fALS subject-derived fibroblasts. (A) Representative images of iMN stained with TMRM and Hoechst, with equalized contrast (EQ) to enhance visualization and segmentation of cell bodies. Scale bar, 250 µm. (B) Quantification of total TMRM intensity per cell, showing increased mitochondrial membrane potential in SOD1^A4V^, C9orf72, and TDP-43^N352S^ and decrease in CHCHD10^R15L^ mutant iMN compared to controls. (C) Mitochondrial count per cell, indicating a significant increase in the number of mitochondria in TDP-43^N352S^ mutant iMN compared to controls. (D) Average area of individual mitochondria, with TDP-43^N352S^ mutant iMN showing reduced mitochondrial size relative to controls. (E) Average TMRM intensity of individual mitochondria, with TDP-43^N352S^ mutant iMN showing increased mitochondrial polarization to controls. For B-E, n=100 cells for each line (7 control, 3 SOD1^A4V^ 3 C9orf72, 4 TDP-43^N352S^, 4 FUS^R521G^, 3 CHCHD10^R15L^). (F) Example kymograph of mitochondrial motility along neurites, illustrating the path and directionality of mitochondrial movement. The upper panel shows a tracked neurite and the lower panel the resulting kymograph and tracked mitochondria. (G) Percentage of motile mitochondria per cell, showing increased motility in TDP-43^N352S^ mutant iMN. (H) Total distance traveled by mitochondria in neurites, with TDP-43^N352S^ mutants showing increased mitochondrial transport distance. (I) Average speed of mitochondrial movementdid not differ significantly across genotypes. Error bars represent SEM, *p ≤ 0.05, **p ≤ 0.01, ***p ≤ 0.001, ****p ≤ 0.0001.

To measure mitochondrial motility in iMN neurites, we labelled cells with TMRM and recorded live imaging time series to generate kymographs along the neurites (Fig. 5F). Compared to controls, TDP-43^N352S^ iMN exhibited mitochondrial motility alterations, including higher percentage of motile mitochondria (Fig. 5G) and higher total distance travelled (Fig. 5H). The average speed (Fig. 5I) and ratio between retro- and anterograde distance travelled was not different between control mitochondria and any of the fALS groups (not shown).

To assess bioenergetic fluxes in iMN, we measured oxygen consumption and extracellular acidification rates (OCR and ECAR, respectively) using the Seahorse bioanalyzer system. We found no difference in basal respiration between control and fALS lines (Fig. 6A). Maximal OCR capacity was elevated in SOD1^A4V^, TDP-43^N352S^, and CHCHD10^R15L^ iMN (Fig. 6B). TDP-43^N352S^ and CHCHD10^R15L^ iMN also exhibited higher rates of oligomycin sensitive (i.e., ATP generating) respiration (Fig. 6C). Spare respiratory capacity (i.e., difference between basal and maximal uncoupled respiration) was increased in SOD1^A4V^ and CHCHD10^R15L^ iMN compared to controls (Fig. 6D). Baseline ECAR, indicative of glycolytic conversion of pyruvate into lactate, was significantly decreased in TDP-43^N352S^ mutants (Fig. 6E), which resulted in increased OCR/ECAR ratio (Fig. 6F), indicating a shift towards oxidative metabolism in these iMN.

**Figure 6.**
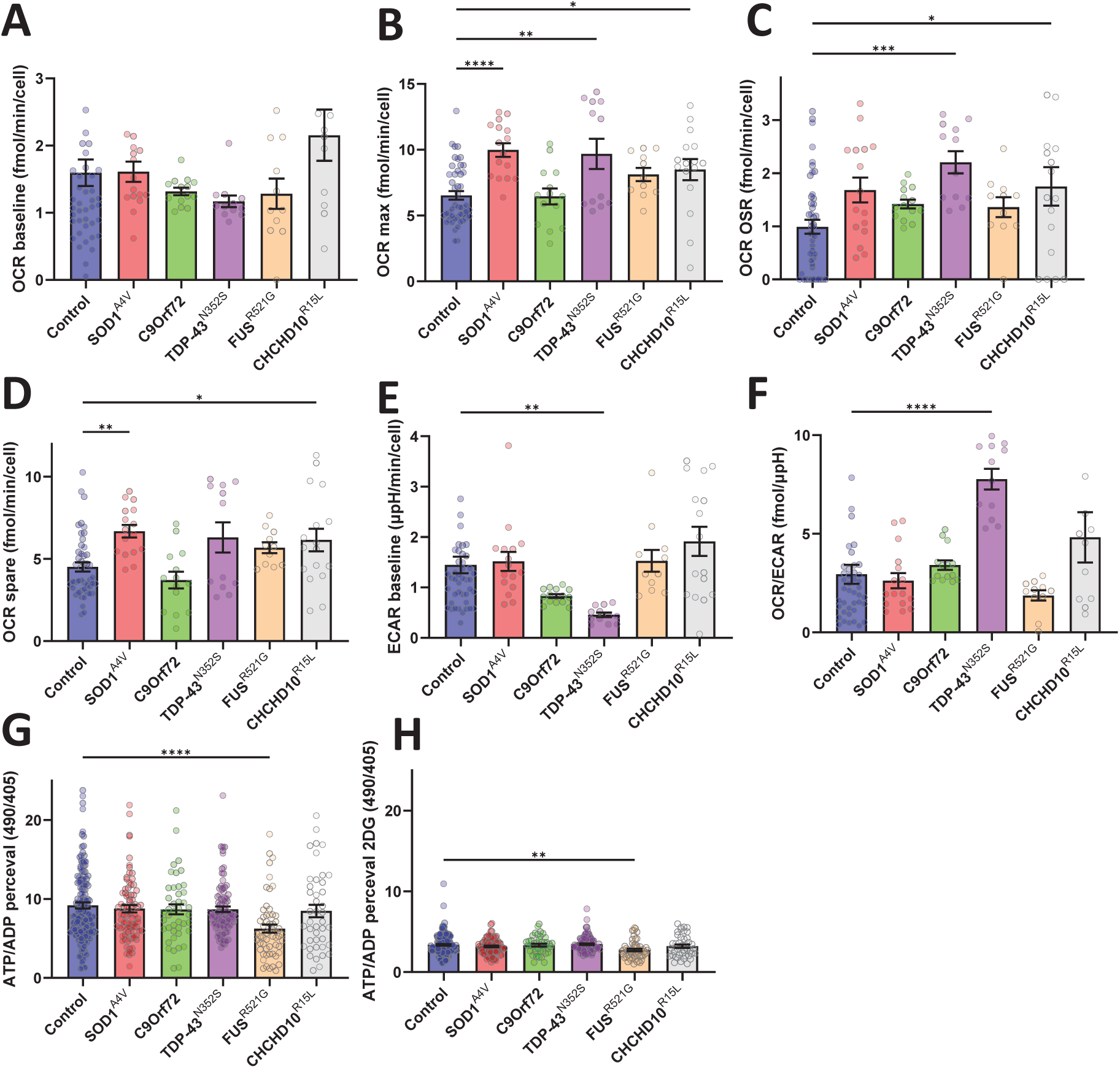
Bioenergetic profile and ATP/ADP ratios in iMN from control and fALS subject-derived fibroblasts. (A) Basal oxygen consumption rate (OCR) across control and fALS iMN, showing no significant differences among groups. (B) Maximal OCR, with significant increases observed in SOD1^A4V^, TDP-43^N352S^, and CHCHD10^R15L^ mutant iMN compared to controls. (C) Oligomycin-sensitive OCR, indicating enhanced mitochondrial respiration in TDP-43^N352S^ and CHCHD10^R15L^ mutants. (D) Spare respiratory capacity, elevated in SOD1^A4V^ and CHCHD10^R15L^ mutants. (E) Baseline extracellular acidification rate (ECAR), significantly decreased in TDP-43^N352S^ iMN. (F) OCR/ECAR ratio, showing an increased oxidative metab iMN olism in TDP-43^N352S^ iMN. (G) Steady-state ATP/ADP ratio, demonstrating a reduction in FUS^R521G^ iMN compared to controls. (H) ATP/ADP ratio following 2-deoxy-D-glucose treatment, highlighting sustained deficits in FUS^R521G^ iMN. *p ≤ 0.05, **p ≤ 0.01, ***p ≤ 0.001, ****p ≤ 0.0001.

Lastly, to assess the energy state of the iMN, we transduced them with lentivirus expressing the biosensor PercevalHR that measures intracellular ATP/ADP ratios. FUS^R521G^ iMN mutants had significantly decreased steady state ATP/ADP ratio in standard culture conditions (Fig. 6G) and after the addition of 2-deoxy-D-glucose to inhibit glucose utilization (Fig. 6H).

Together, these results indicate that fALS iMN display alterations of mitochondrial and bioenergetic functions including, mitochondrial polarization, mitochondrial trafficking, and cellular ATP/ADP levels. These alterations, however, are quite different among fALS mutations, with TDP-43^N352S^ iMN exhibiting changes in almost all the parameters investigated. Moreover, these findings suggest that the widespread bioenergetic and mitochondrial alterations do not always correspond to energy production defects but may instead represent adaptive changes to maintain homeostasis.

### Mutation-specific fALS iMN survival defects are improved by small molecules targeting energy metabolism and oxidative stress

To measure survival of iMN we developed an automated single cell iMN detection and tracking pipeline (Fig. 7A). iMN cultures were infected with hSyn-eGFP on D8 and imaged on every other day from D12 to D46 using the pipeline described above (Fig. 3A). Training data was generated manually to teach a machine learning algorithm to recognize and track individual cells exhibiting neuronal morphology, similar to an approach previously published for acute tracking of iPSC derived neurons^21^. An example is shown in Fig. 7B, where the orange rectangles indicate an individual iMN marked as ‘cell’ with an assigned identity number. This approach removes artifacts resulting from different transformation rates and cell densities between lines and batches, which could arise with endpoint counting approaches. For each iMN, the starting point for Kaplan-Meier analysis was the time when the cell was recognized by the AI as an iMN, and the endpoint was when the cell disappeared from the culture, indicating death of the cell. Lifelines were censored after the last timepoint (D46). Compared to controls, SOD1^A4V^, C9orf72, and FUS^R521G^ mutant iMN had a significantly higher risk of death (Fig. 7C). Importantly, parental fibroblast passage number or subject age at time of skin biopsy were not correlated with iMN survival (Supplementary Fig. S2C-D).

**Figure 7.**
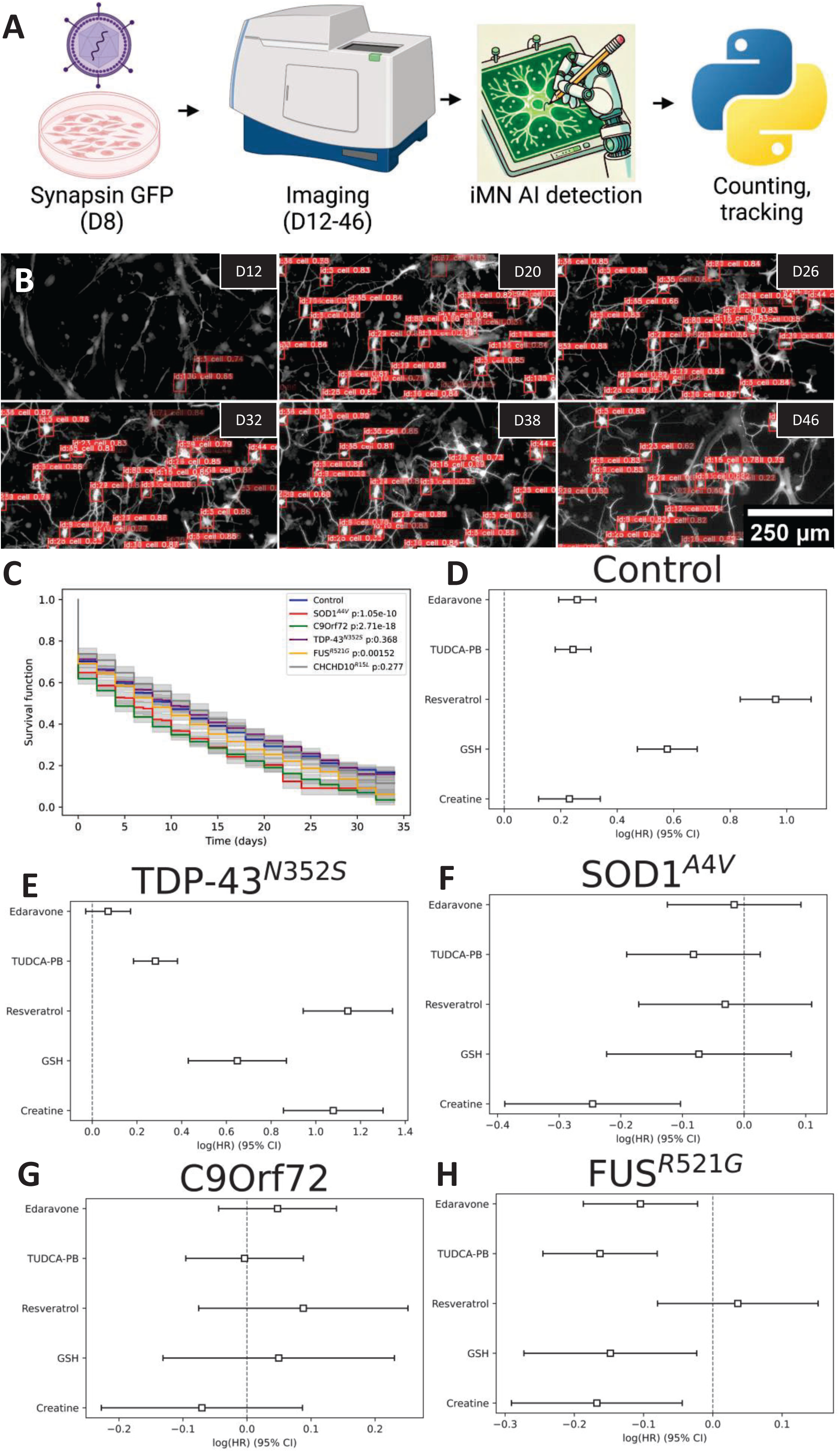
Survival analysis of fALS iMN and effects of therapeutic compounds. (A) Workflow of survival tracking in iMN. Cultures were transfected with Synapsin-GFP on Day 8 (D8) and imaged from Day 12 (D12) to Day 46 (D46). iMN detection and tracking were performed with a finetuned ML model (YOLOv8) to monitor individual cells. (B) Representative images of iMN cultures with tracked cells (identified by orange labels) over time, from D12 to D46. (C) Kaplan-Meier fitted survival from control and fALS iMN under baseline conditions, showing reduced survival in certain fALS lines (p-values are in the legend). (D-H) Cox proportional hazards model hazard ratios of iMN survival for control and fALS mutants under different drug treatments, relative to vehicle (log(HR)=0). Squares represent the log(HR), error bars the 95% confidence interval. n=100-3500 cells for each group (sampled from 7 control, 3 SOD1^A4V^ 3 C9orf72, 4 TDP-43^N352S^, 4 FUS^R521G^, 3 CHCHD10^R15L^ lines, and 1-9 technical replicates from 3 batches of differentiations).

To determine if mitochondrial alterations and oxidative stress may play a role in decreased viability of mutant iMN, we tested the effects of small molecules that target mitochondrial dysfunction, bioenergetics, excitotoxicity, and oxidative stress and have been used in clinical trials of sporadic ALS. Drugs included the FDA approved edaravone, the combination of tauroursodiol with phenylbutyrate (TUDCA-PB), resveratrol, glutathione (GSH), and creatine. Surprisingly, all drugs increased risk for control lines (Fig. 7D) and TDP-43 ^N352S^ iMN (except edaravone that had no effect) (Fig. 7E). Notably, TDP-43 ^N352S^ iMN did not have viability defects in vehicle medium compared to controls (Fig. 7C), possibly because mitochondrial changes were adaptive in nature and the drugs disrupts homeostasis. SOD1^A4V^ mutants had significantly decreased hazard ratios (HR) with creatine (Fig. 7F), whereas for C9orf72 we found no evidence of a beneficial effect with any of the drugs tested (Fig. 7G). Interestingly, FUS^R521G^ iMN responded well to all drugs, with diminished HR for death in comparison to vehicle, except for resveratrol (Fig. 7H). These findings indicate that the viability response to mitochondrial and antioxidant drugs, some of which (riluzole and edaravone) are FDA approved, differs among fALS groups and suggest that similar differences may exist in the much broader and diverse population of sporadic ALS.

## Discussion

In this study, we generated iMN from fibroblasts of both control individuals and individuals with fALS using a DC technique. We found that both control and fALS DC iMN express MN-specific markers and exhibit neuronal morphological features and a large majority of neuronal cells expressed MN markers. Importantly, we also determined that in our hands there were significant line-specific differences in the efficiency of conversion of fibroblasts to neuronal cells. These differences could be quite large and were not associated with passage number or age at skin biopsy and did not appear to be affected by the disease state (i.e., control and fALS). This variability in differentiation efficiency could be due to differences in cellular responses to reprogramming or in the lentiviral infectability, among fibroblast lines and viral batches.

Potential differences in transformation must be considered, especially when estimating the number of independent fibroblast lines to use and planning experiments that involve bulk assays. Here, with the exception of Seahorse flux analyses, for the characterization of iMN phenotypes we opted for studies at the single cell level, as all lines were converted into iMN with sufficient efficacy for this type of analyses. We adapted deep learning frameworks to automate iMN detection under a variety of imaging and experimental conditions, allowing for data collection and evaluation of phenotypes at the single-cell level in an unbiased and time efficient manner.

Our initial characterization of iMN morphology revealed only a few differences among lines, whereby SOD1^A4V^ and FUS^R521G^ iMN had decreased total neurite length or branching. Moreover, as expected, we noted that TDP-43^N352S^ iMN had increased cytoplasmic localization of TDP-43, but in the absence of cytoplasmic inclusions. TDP-43 mislocalization was also present in FUS^R521G^ iMN, which to our knowledge has not previously been described in cellular models. TDP-43 mislocalization is uncharacteristic of SOD1 and CHCHD10 mutants^22,23^, in agreement with our findings. However, while increased TDP-43 cytoplasmic localization and inclusions are present in postmortem tissue of C9orf72 mutation carrying patients^24,25^, we did not find this feature in DC iMN. These finding indicates that DC iMN, at least in our experience, do not recapitulate some key molecular phenotypes of ALS. This finding was in agreement with the lack of TDP-43 proteinopathy reported in unperturbed C9Orf72 iPSC derived iMN^24,26^.

Mitochondrial dysfunction and abnormal bioenergetics have been associated with both sporadic ALS and fALS, but these alterations have not previously been investigated in DC iMN. We observed increased mitochondrial membrane potential in SOD1^A4V^, C9orf72, and TDP-43^N352S^ iMN, indicating increased mitochondrial function (Supplementary figure S4 summarizes the mitochondrial phenotypes). In TDP-43^N352S^ iMN, individual mitochondria were more polarized, and there were changes in mitochondrial morphology and increased mitochondrial motility in neurites. Furthermore, SOD1^A4V^, TDP-43^N352S^, and CHCHD10^R15L^ mutant iMN exhibited increased maximal OCR and ECAR). These changes could result from enhanced bioenergetic activity in fALS iMN, potentially a hypermetabolic state like that observed in ALS subjects and sporadic ALS fibroblasts^15^. However, when we measured ATP/ADP ratios with the genetically encoded biosensor PercevalHR, most fALS mutants were not significantly different than controls, and only FUS^R521G^ iMN had decreased ATP/ADP ratio. A hypermetabolic state with unchanged ATP/ADP ratio may indicate an adaptive response to increased energy demands in fALS iMN. Hypermetabolism and mitochondrial hyperpolarization could cause increased reactive oxygen species generation, which requires upregulation of antioxidant defenses. contributes to motor neuron degeneration. In FUS^R521G^ iMN, there was no hypermetabolism, as previously shown in iPSC-derived FUS mutant iMN^27^. In these cells, the lack of metabolic adaptation may cause a lower ATP/ADP ratio.

To follow the fate of individual iMN longitudinally, we developed a technique to track single iMN in culture and assess their lifespan. We were able to follow individual cells as they differentiated and aged in the dish, until their death which typically occurred within a few weeks from the time when they differentiated into identifiable iMN. Whereas TDP-43^N352S^ and CHCHD10^R15L^ iMN did not differ from controls, SOD1^A4V^, C9orf72, and FUS^R521G^ iMN exhibited higher risk of death. We hypothesized that the hypermetabolic phenotypes may be adaptive in TDP-43^N352S^ and CHCHD10^R15L^ iMN to fulfil energy demands, but in SOD1^A4V^, C9orf72, and FUS^R521G^ iMN the metabolic changes could be insufficient or detrimental. To begin testing this hypothesis, we treated cells with small molecules that have been proposed to alleviate cellular and metabolic stress. These included edaravone, a redox-scavenging molecule that is FDA approved for ALS^28–30^, the combination of TUDCA and PB, which targets ER stress and Bax-driven mitochondrial cytochrome c release^31,32^, resveratrol, a SIRT1 activator that enhances mitochondrial biogenesis and quality control^33,34^, glutathione, an essential antioxidant, and creatine, a high-energy phosphate storage to replenish ATP in cells^35^. Surprisingly, control and TDP-43^N352S^ iMN survival was negatively affected by most treatments (except edaravone for TDP-43^N352S^), suggesting that these lines were in a homeostatic state that was disrupted by the drugs. None of the treatments decreased the death risk of C9orf72 iMN, whereas creatine improved survival in SOD1^A4V^ and FUS^R521G^ iMN. FUS^R521G^, the only fALS iMN group with ATP/ADP defects, benefited from creatine, edaravone, TUDCA-PB, and GSH.

Taken together, these results suggest that the energetic phenotypes of DC iMN with different fALS mutations could be linked to their survival outcomes. SOD1^A4V^, C9orf72 repeat expansions, and FUS^R521G^ iMN exhibit bioenergetic hyperactivity and higher risk of cell death, indicating that their hypermetabolic state may be detrimental, leading to accelerated degeneration. In FUS^R521G^ iMN, we found evidence of bioenergetic incompetence and creatine, improved their survival. FUS^R521G^ iMN also benefitted from antioxidants and TUDCA plus PB, suggesting that they may also undergo increased oxidative stress.

The variable responses to the treatments in the different fALS iMN highlight the importance of identifying adequate model systems for molecule screening, as different fALS mutations may require specific interventions to effectively mitigate neurodegeneration. For these reasons, the DC fALS iMN system used in this study, although not well suited for large drug screens, may be developed into a targeted platform, where candidate drugs could be tested for each mutation group or even for each subject within a few weeks, if fibroblasts are available.

The image analysis pipeline that we have developed using machine learning to identify iMN and measure mitochondrial morphological and functional parameters as well as the strategy used to identify and follow individual iMN longitudinally in culture may prove to be useful tools for future of larger cohorts of DC iMN derived from subjects affected by ALS. For example, the combination of optimized DC iMN protocols with these automated analysis tools could be implemented in the future for mechanistic studies and drug testing using DC iMN from sporadic ALS fibroblasts, and possibly other neurodegenerative diseases where neuronal mitochondrial abnormalities have also been described.

There are of course limitations to the iMN modeling approach, some of which could be improved while others are inherent to the system and will be difficult to address. The former include the variable differentiation efficacy, which can likely be improved by further optimization of the gene delivery and selection systems. Moreover, the lifespan of DC iMN is relatively short compared, for example, to iPSC-derived iMN, but it is possible that optimization of media composition and growth conditions may increase the lifespan of the cells. The latter involve the natural senescence of the parental fibroblasts. To partially mitigate this problem, we typically maintain several frozen vials of early passage fibroblasts for DC iMN generation, but this is not an unlimited source of parental cells. It remains to be determined whether genetic modifications, such as telomerase expression, could be implemented to prolong the viability of cultured fibroblasts for DC iMN.

## Acknowledgements

This work was supported by Muscular Dystrophy Association Development Grants MDA602762 (to CK) and MDA961871 (to HK), Project ALS/NextGen ALS grant ALS215291 (to HK and GM), and NIH/NINDS grants R35NS122209 (to GM), R01NS139141 (to HK).

This study used an anonymized fibroblast line from the NINDS Repository, as well as clinical data related to the cell line. NINDS Repository sample number corresponding to the sample used is (ND39022). Other lines used in this study were shared by the groups of Dr. John Ravits (UCSD), Haining Zhu (U of A), Hiroshi Mitsumoto (Columbia U), and Neil Shneider (Columbia U).

## Supplemenrary figure legends

Supplementary Figure 1. Performance of the Resnet50 architecture machine learning model for iMN classification. (A) Accuracy plot showing training (blue) and test (red) accuracies over 10 epochs for the model used to classify cells as dead, non-neuron, or iMN. (B) Receiver Operating Characteristic (ROC) curve with Area Under the Curve (AUC) values for each class on the holdout test set, demonstrating the model’s performance in distinguishing between dead, non-neuron, and iMN classes. (C) Confusion matrix for the classification model on the holdout test set, showing the true and predicted labels for each category. (D) Normalized confusion matrix, displaying the fraction of correctly and incorrectly classified labels for dead, non-neuron, and iMN categories. (E-G) Example images of cells in each classification category—dead, non-neuron, and iMN—from three different staining conditions: hSyn-eGFP live imaging (E), TDP-43 immunocytochemistry (F), and TMRM live imaging (G). Red arrowheads indicate the cell (or debree) in the exact center which is being classified.

Supplementary Figure 2. Training and validation metrics, model precision-recall, and regression analyses for iMN classification and survival analysis. (A) Training and validation loss curves for box loss, classification loss, distance focal loss (dfl), Mean Average Precision (mAP) across different Intersection over Union (IoU) thresholds for bounding box detection, and Precision-recall curve, with the average precision across all classes and specific mAP scores at different IoU thresholds during model training, showing the convergence of these metrics over 6000 steps. Precision - recall metrics for the model on the validation set (B).

Supplementary Figure 3. iMN survival half-life (A,B,C) and transformation efficiency (D, E, F) condounded by passage (A, D), age at biopsy (B, E), and sex (C, F), showing R-squared values and p-values for each relationship, suggesting no confounding effects in our data.

Supplementary Figure 4. Summary of key bioenergetic findings from iMN analyses across control and fALS genotypes. Heatmap displaying percent changes for multiple mitochondrial and bioenergetic parameters in iMN derived from fALS versus control iMN. Percent changes are calculated relative to control iMN, with minus log2-transformed p-values indicated in parentheses. Only parameters with statisticaly significant differences are shown.

