## Supplementary figures and images for "Investigation of mitochondrial phenotypes in motor neurons derived by direct conversion of fibroblasts from familial ALS subjects"

### Supplementary Figures

#### Supplementary Figure 1

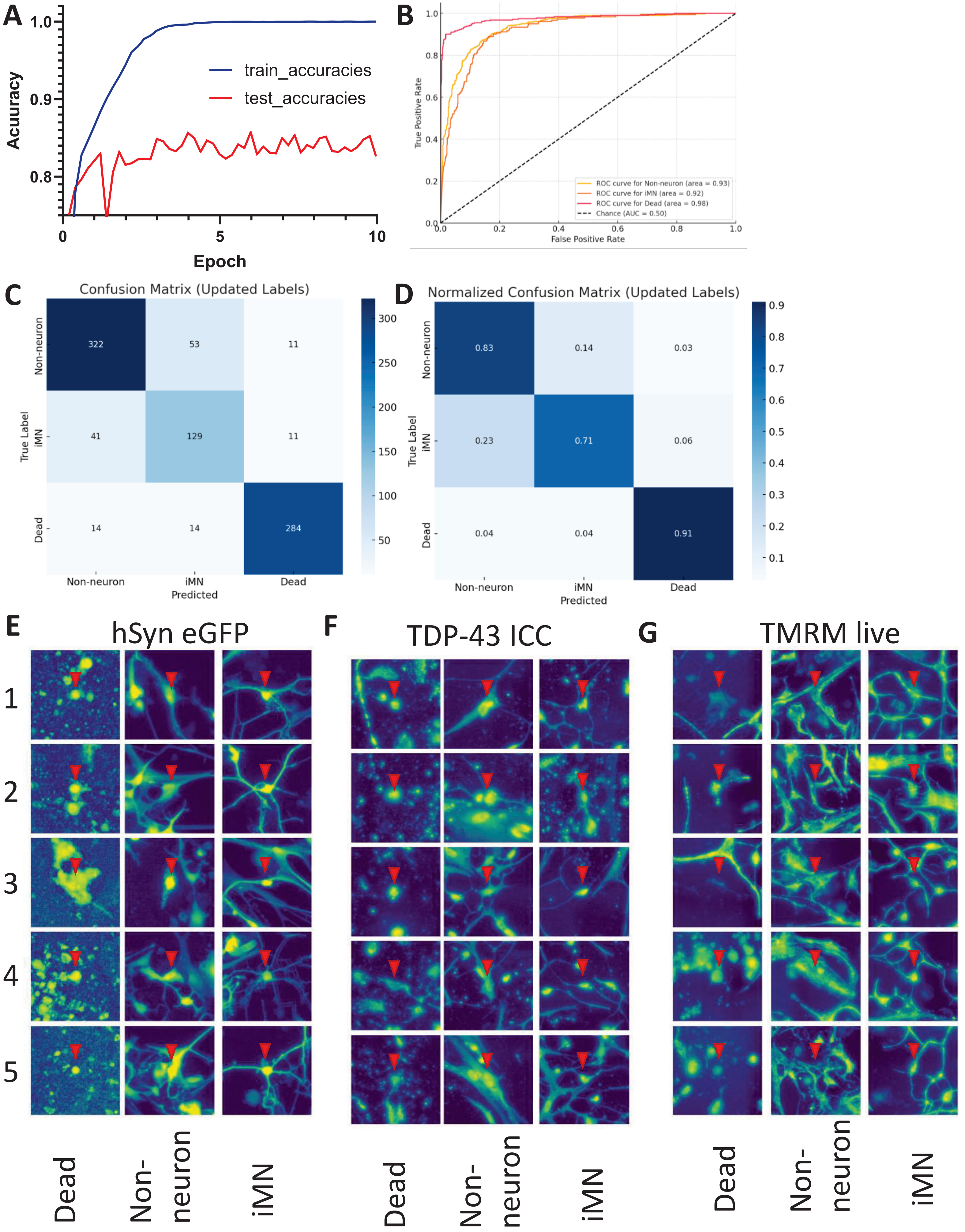

#### Supplementary Figure 2

**A**

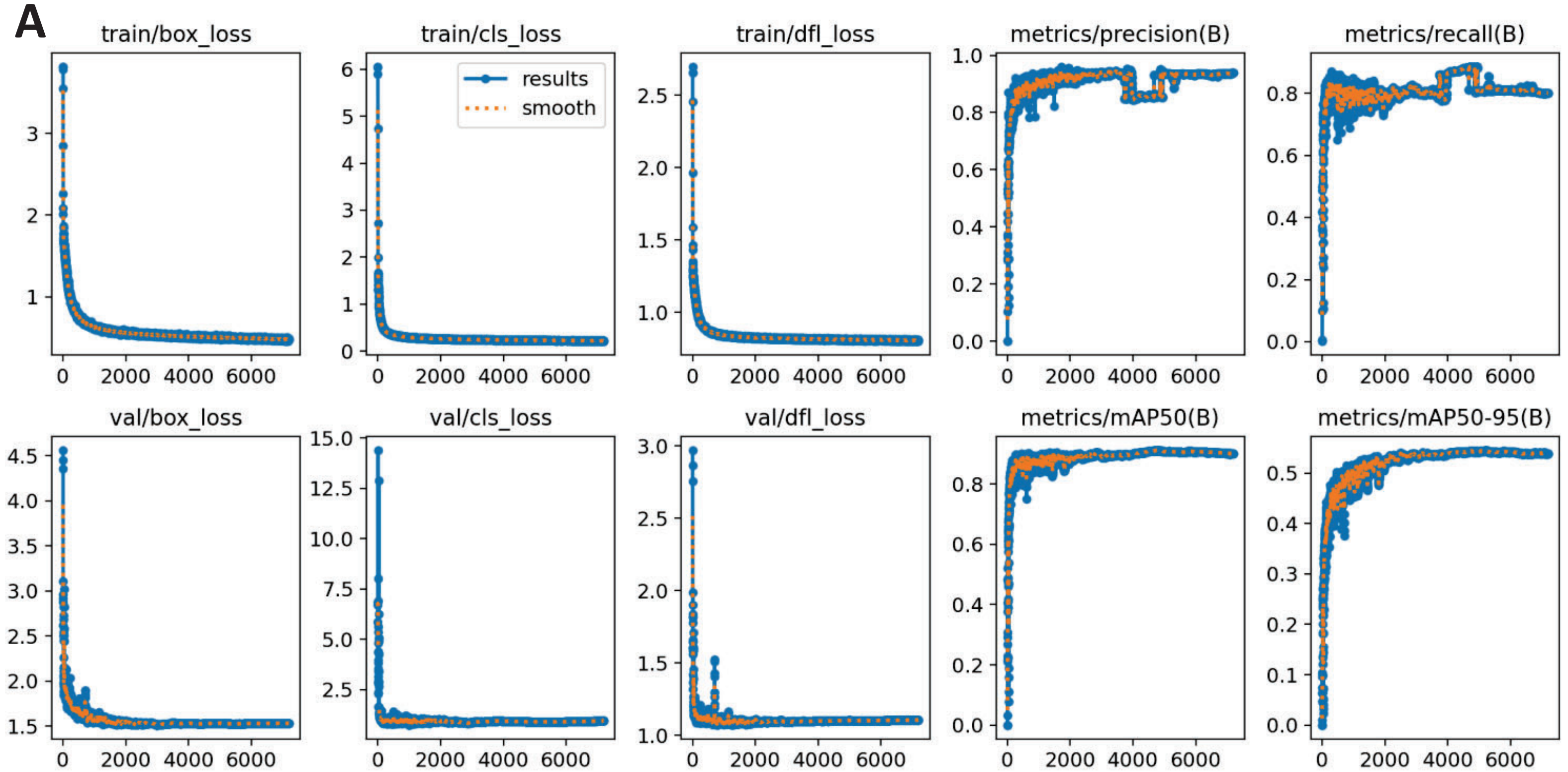


**B**

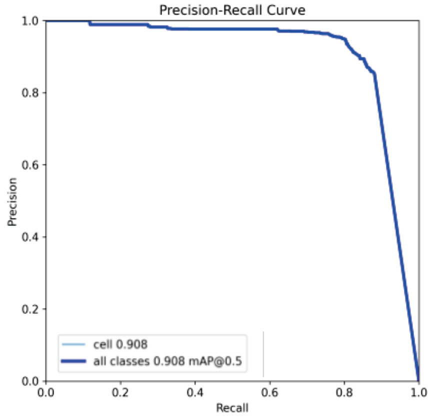

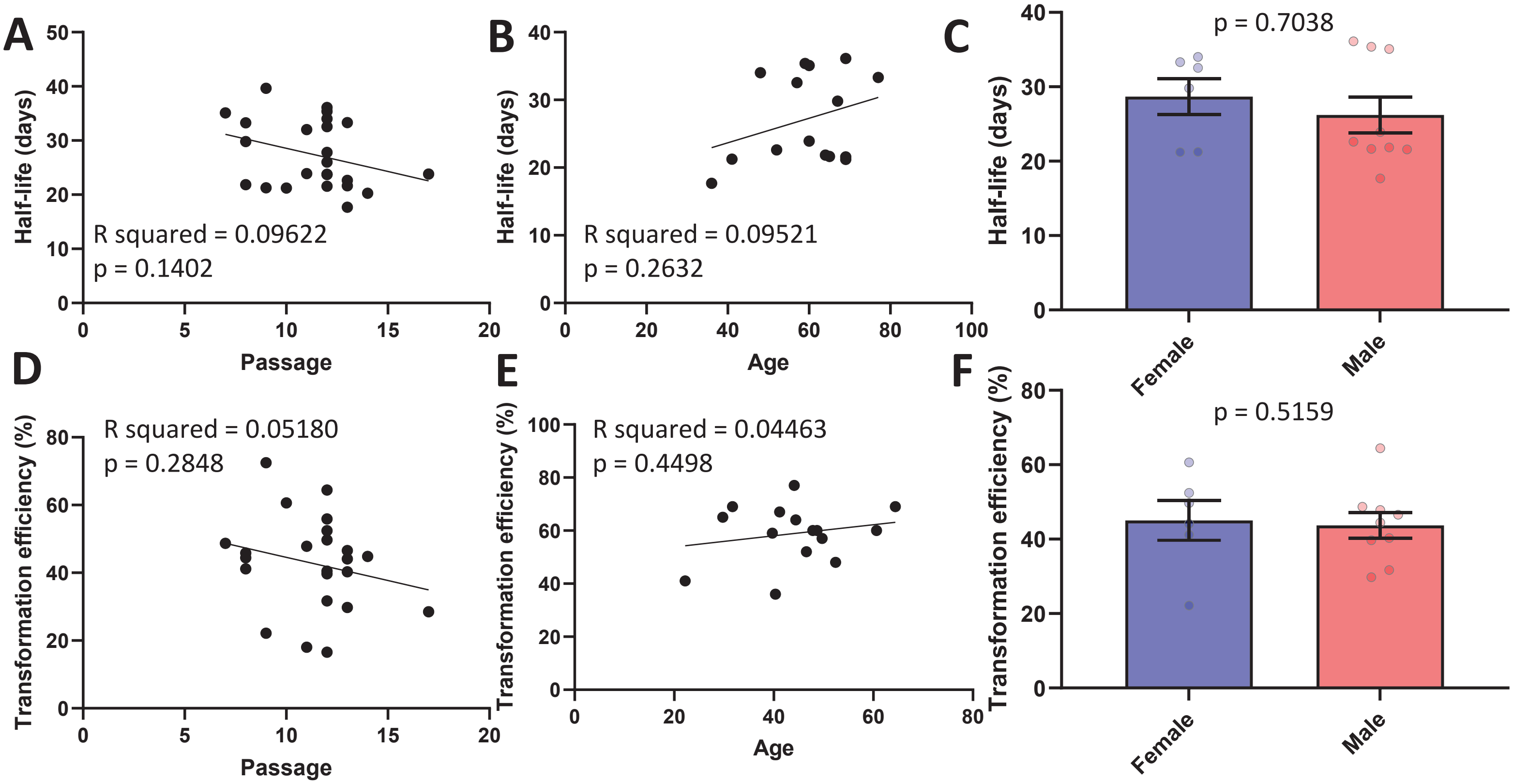

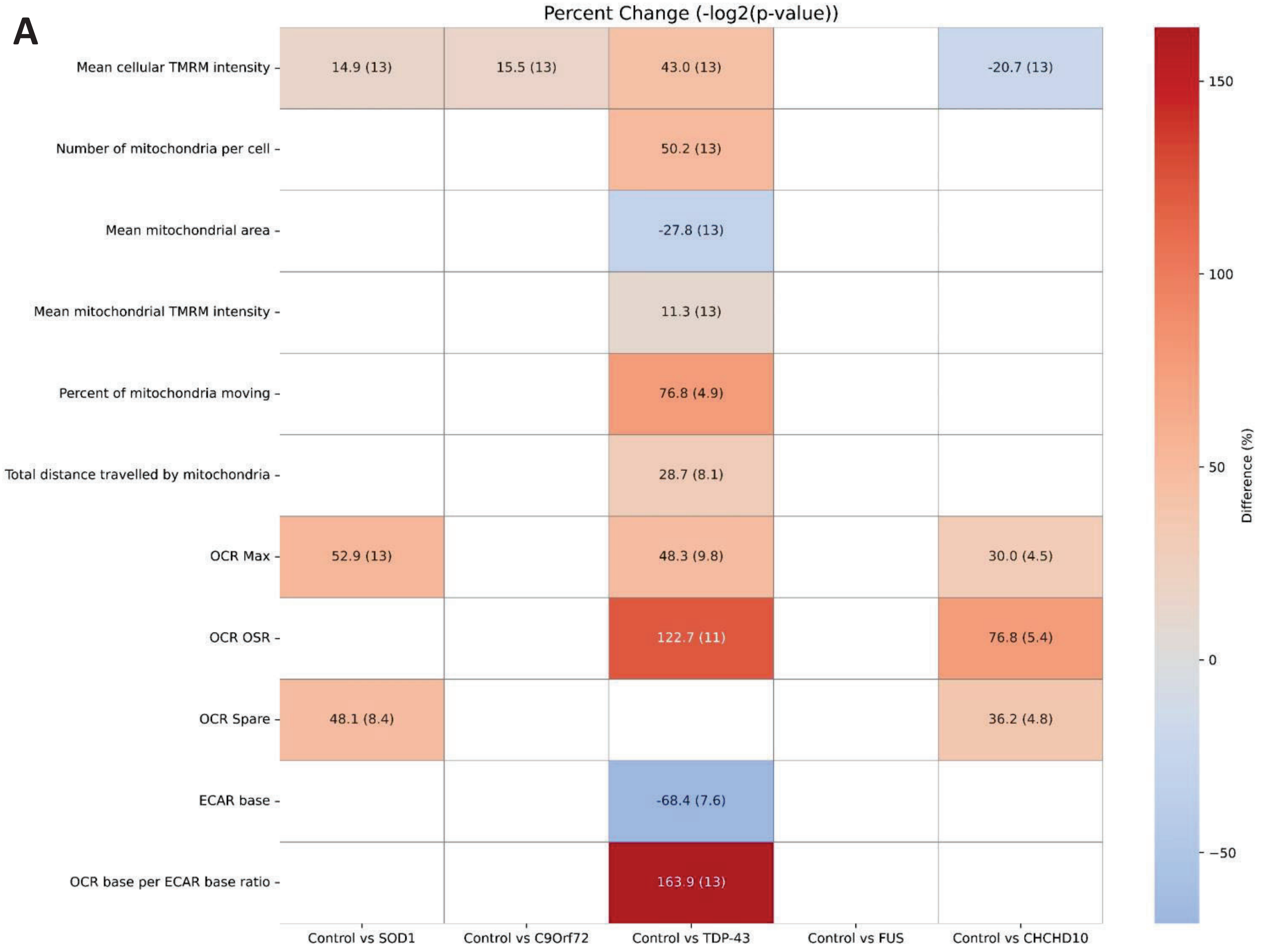
